# Secondary metabolites produced by *Colletotrichum* spp. on different olive cultivars

**DOI:** 10.1101/2022.11.18.517023

**Authors:** Mario Riolo, Carlos Luz, Elena Santilli, Giuseppe Meca, Santa Olga Cacciola

## Abstract

This study was aimed to characterize the secondary metabolites produced by four *Colletotrichum* species, *C. acutatum*, *C. gloeosporioides*, *C. godetiae* and *C. karsti*, both *in vitro*, on potato dextrose agar (PDA) and oatmeal agar (OA), and during the infection process of fruits of four olive cultivars differing in susceptibility to anthracnose, ‘Coratina’ and ‘Ottobratica’, both susceptible, ‘Frantoio’ and ‘Leccino’, both resistant. The metabolites were extracted from axenic cultures after seven days incubation and from olives at three different times, 1, 3 and 7 days post inoculation (dpi). They were identified using the HPLC-QTOF analysis method. In total, as many as 45 diverse metabolites were identified; of these 29 were detected on infected olives and 26 in axenic cultures on agar media (OA and PDA). Only 10 metabolites were present in both fruits and axenic cultures while 19 were found exclusively on olives and 16 exclusively in axenic cultures. The 45 identified metabolites comprised fatty acid, phenolics, pyrones, sterols, terpenes and miscellaneous compounds. Each *Colletotrichum* species produced a different spectrum of metabolites depending of the type of matrices. On artificially inoculated olives the severity of symptoms, the amount of fungal secondary metabolites and their number peaked 7 dpi irrespective of the cultivar susceptibility and the virulence of the *Colletotrichum* species. The metabolite profiles as represented by heat maps were the result of the interaction olive cultivar x *Colletotrichum* species.

## 1. Introduction

Anthracnose is a major disease of olive (*Olea europaea*) in many olive growing countries worldwide (Cacciola et al., 2012; Kolainis et al., 2020; Moral et al., 2021; Talhinhas et al., 2018). Several species of *Colletotrichum* are responsible for olive anthracnose (OA). Overall, 18 *Colletotrichum* species have been so far reported to be associated with this disease worldwide (Talhinhas et al., 2018); they vary in virulence and geographical distribution (Moral et al., 2021; Moreira et al., 2021; Riolo et al., 2022; Schena et al., 2014; Talhinhas et al., 2018). Most of these species were segregated from the *Colletotrichum acutatum* (*C. acutatum s.l*.), *C. boninense* (*C. boninense s.l*.) and *C. gloeosporioides* (*C. gloeosporioides s.l*.) species complexes and formally described as separate taxa after the acceptance of multilocus phylogenetic analysis as a taxonomic criterion. *Colletotrichum acutatum* is the most virulent species; as a pathogen responsible for OA it is widespread in the southern hemisphere and is emerging in several olive growing areas of the Mediterranean macroregion, including Albania, Greece, Italy, Portugal, Spain and Tunisia (Riolo et al., 2022; Schena et al., 2017). *Colletotrichum nymphaeae* is the dominant *Colletotrichum* species associated with OA in south-western Portugal and was reported as the second most common species associated with OA, after *C. acutatum*, in Uruguay (Moreira et al., 2021; Riolo et al., 2022; Talhinhas et al., 2015). It has been found also in other olive producing areas of the Mediterranean macroregion but has not yet been reported from olive producing areas of the southern hemisphere, except for Brazil (Moral et al., 2021). *Colletotrichum godetiae*, along with *C. acutatum* is the most common species associated with OA in Greece, Southern Italy and Tunisia. It has been recorded also in Montenegro with the former name of *C. clavatum* and is by far the dominant species associated with OA in Andalusia, the leading olive producing region of Spain (Faedda et al., 2011; Moral et al., 2021; Riolo et al., 2022). Other *Colletotrichum* species are weakly pathogenic, occur sporadically or have a more restricted geographical distribution (Moral et al., 2021, 2015; Riolo et al., 2022; Schena et al., 2014). *Colletotrichum fioriniae* was first reported from some regions of Portugal among the *Colletotrichum* species associated with OA (Cacciola et al., 2012) and subsequently as the species responsible for OA outbreaks in Uruguay and California (USA) (Moral et al., 2021; Moreira et al., 2021). In the latter country it is also common as causal agent of anthracnose of pistachio (*Pistacia vera*) (Moral et al., 2021). Very recently this *Colletotrichu*m species has been reported from southern Italy as causal agent of olive anthracnose in association with *C. acutatum* and *C. godetiae* (Riolo and Cacciola, 2022). *Colletotrichum karsti*, like other *Colletotrichum* species associated with OA, is polyphagous (Talhinhas and Baroncelli, 2021) but it was proved to be a weak pathogen on olive (Riolo et al., 2022; Schena et al., 2014). *Colletotrichum siamense* was found to be associated with OA in Australia and, differently from *C. karsti*, in artificial inoculations on detached olives proved to be aggressive (Moral et al., 2021; Schena et al., 2014). *Colletotrichum theobromicola*, alone or in association with other *Colletotrichum* species, was reported as causal agent of anthracnose in Australia and South America (Lima et al., 2020; Moreira et al., 2021; Schena et al., 2014), and in artificial inoculations on detached olives proved to be very aggressive (Schena et al., 2014). It has not been reported so far as a pathogen of olive in the northern hemisphere but has been intercepted in Israel as causal agent of fruit anthracnose on avocado (*Persea americana*) and leaf spot on cyclamen (*Cyclamen persicum*) (Sharma et al., 2017, 2016). Recently, *C. theobromicola*, along with *C. aenigma*, *C. alienum*, *C. perseae* and *C. siamense*, all already known as pathogens of olive (Moral et al., 2021; Schena et al., 2014), have been recognized as noxious organisms of quarantine concern for the European Union (EFSA Journal 2022). The most typical and economic relevant symptom of OA is fruit rot, which results in yield losses due to premature fruit drop and in detrimental effects on physicochemical and organoleptic characteristics of the oil produced with infected fruits (Cacciola et al., 2012; Gouvinhas et al., 2019; Graniti et al., 1993; Kolainis et al., 2020; Leoni et al., 2018; Moral et al., 2021, 2014; Peres et al., 2021; Romero et al., 2022; Runcio et al., 2008; Talhinhas et al., 2018). The severity of damage caused by OA depends on several factors, including environmental conditions, the virulence of the *Colletotrichum* species involved and the susceptibility of olive cultivar. A great variability in susceptibility to OA among olive cultivars of various geographic origin has been observed (Cacciola et al., 2012; Moral et al., 2017, 2015, 2008; Moral and Trapero, 2009; Talhinhas et al., 2018, 2015). Among the Italian cultivars ‘Frantoio’ and ‘Leccino’ were proved to be relatively resistant while ‘Carolea’, ‘Coratina’ and ‘Ottobratica’ were proved to be very susceptible (Riolo et al., 2022). Moreover, a significant interaction between olive cultivar susceptibility and *Colletotrichum* species virulence was highlighted (Riolo et al., 2022; Talhinhas et al., 2015). For example, ‘Picual’, a Spanish olive cultivar which is regarded as relatively resistant in the geographical area of origin where *C. godetiae* has been largely prevailing over other *Colletotrichum* species for many years (Cacciola et al., 2012; Moral et al., 2021), was confirmed to be resistant to *C. godetiae* but proved to be very susceptible to *C. acutatum* in laboratory tests on detached fruits (Riolo et al., 2022).

In addition to fruit rot, symptoms of OA include blossom blight, leaf yellowing, wilting defoliation and dieback of twigs and branches (Cacciola et al., 2012; Moral and Trapero, 2009; Moreira et al., 2021). The very low frequency of pathogen isolation from leaves and branches of adult trees with natural infections of OA, suggested the hypothesis that a single or multiple phytotoxins produced by *Colletotrichum*, could account for the induction of these symptoms on leaves and twigs (as reviewed by Cacciola et al. (Cacciola et al., 2012)). This hypothesis was corroborated by Moral et al. (Moral and Trapero, 2009). These Authors inoculated young olive plants and noticed that wilting and dieback symptoms developed only on twigs bearing rotten fruits. Twigs without fruits were also infected, but did not show symptoms, suggesting that a diffusible metabolite produced by the fungus in infected fruits is responsible for the symptoms on other organs of the plant (Moral et al., 2021). Indeed, Ballio et al. (Ballio et al., 1969) had demonstrated that the causal agent of OA produced *in vitro* aspergillomarasmin B, a phytotoxic derivative of lycomarasmin. Numerous other bioactive secondary metabolites produced *in vitro* by *Colletotrichum* species have been since isolated and identified from culture fluids, they mostly included polyketides, terpenoids, peptides and aminoacid derivatives, alkaloids, derivatives of shikimic acid and metabolites of mixed biosynthetic origin (Chakraborty and Ray, 2021; Chapla et al., 2014; Chen et al., 2019; García-Pajón and Collado, 2003; Kim and Shim, 2019; Li et al., 2022; Masi et al., 2020; Moraga et al., 2019; Sun et al., 2022; Xu et al., 2019, 2021). The interest of above mentioned studies has focused mainly on the potential applications of these metabolites either in pharmacology as drugs or in agriculture as natural substances with antifungal and herbicidal activity (Kim and Shim, 2019; Masi et al., 2017). Several secondary metabolites isolated from culture fluids of *Colletotrichum* species, such as Colletotrichin E, Colletofragarone A1, Colletochlorin A and Aspergillomarasmin showed phytotoxic activity (Chapla et al., 2014; Masi et al., 2017). However, after the pioneering study of Ballio et al. (Ballio et al., 1969) no other research has dealt with the characterization of secondary metabolites produced by *Colletotrichum* species infecting olive fruits, their detrimental effects on the oil quality and their role as pathogenicity or virulence factors in the pathogenesis of OA. This study, aimed at identifying the secondary metabolites produced by diverse *Colletotrichum* species during the infection process of olive fruits, was intended as a contribute to fill this gap. In particular, its aim was to identify the secondary metabolites produced by diverse *Colletotrichum* species in vitro and during the infection process of olive fruits.

## 2. Materials and Methods

### 2.1 Fungal isolates, culture media and production of inoculum

Isolates of four different *Colletotrichum* species, *C. acutatum*, *C. gloeosporioides*, *C. godetiae* and *C. karsti*, were included in this study (Table 1). The isolates were Identified in previous studies based on both morphological characteristics and phylogenetic analysis of the internal transcribed spacer (ITS) ITS1–5.8S–ITS2 region of the ribosomal DNA (rDNA), and part of the β-tubulin gene (TUB2) sequences (Cacciola et al., 2020; Riolo et al., 2022).

**Table 1.** Isolates of *Colletotrichum* used in this study.

| Isolate code | Species | Host<br>(species and cultivar) | Organ | Geographical origin | GenBank Accession |  |
| --- | --- | --- | --- | --- | --- | --- |
| | | | | | ITS-rDNA | β-tubulin 2 |
| UWS149 | *C. acutatum* | *Olea europaea* | Fruit | Australia | MT997785 | MW001517 |
| C9D2C | *C. acutatum* | *O. europaea* | Fruit | Calabria (IT) | MZ502315 | MZ508448 |
| C2 | *C. gloeosporioides* | *Citrus xlimon* | Fruit | Calabria (IT) | JN121211 | JN121298 |
| RD9B | *C. gloeosporioides* | *O. europaea* | Fruit | Calabria (IT) | MZ502314 | MZ508447 |
| OLP 12 | *C. godetiae* | *O. europaea* | Fruit | Calabria (IT) | JN121131 | JN121218 |
| OLP 16 | *C. godetiae* | *O. europaea* | Fruit | Calabria (IT) | JN121137 | JN121224 |
| C12D1A | *C. karsti* | *O. europaea* | Fruit | Sicily (IT) | ON231821 | ON246203 |
| CAM | *C. karsti* | *Camellia* sp. | Leaf | Sicily (IT) | KC425664 | KC425716 |

All isolates included in this study were sourced from the culture collection of the Molecular Plant Pathology Laboratory of the Department of Agriculture, Food and Environment of the University of Catania, Italy. Potato dextrose agar (PDA; Oxoid Ltd., Basingstoke, UK) and oatmeal agar (OA; Sigma-Aldrich, St. Louis, Mo, USA), were used as culture media throughout this study.

A conidial suspension of each isolate (10^6^ conidia ml^-1^) in sterile distilled water (SDW) was prepared from 10-day old cultures grown on PDA at 25 °C and used as inoculum. The concentration of conidia in the suspension was adjusted with a hemocytometer.

### 2.2 Inoculations of olives

Fruits of four olive (*Olea europaea* L.) cultivars (‘Coratina’, ‘Frantoio’, ‘Leccino’ and Ottobratica’) differing in their susceptibility to OA were used in this study; they were collected from 15-year-old olive trees in the experimental orchard of CREA OFA (Council for Agricultural Research and Economics, Research Centre for Olive, Fruit and Citrus Crops) in the municipality of Rende, province of Cosenza, Calabria region, Italy (DATUM WGS 84, 39° 21’ 59.4” N 16° 13’ 44.4” E). The ripening stage of fruits was evaluated according to the maturity index (MI) of Guzmán et al. (Guzmán et al., 2015), with value 0 corresponding to >50% bright green and value 4 to 100% blackish-purple or black. Fruits of the same cultivar were collected from three distinct trees at a medium ripening stage (MI 3-4) and pooled together. They were stored for 3 h inside a refrigerated bag, before being transported to the laboratory and were inoculated the day after collection. Before the inoculation, fruits were surface disinfected by immersion in a 0.5% NaOCl solution for 30 s, rinsed in SDW, blotted dry and placed in incubation trays. Olives were punctured with a sterile needle in an equatorial position and a 20 μl droplet of the conidial suspension (10^6^ conidia/ml), prepared as described above, was pipetted onto the surface of the wound. Controls received a 20 μl droplet of SDW. Forty fruits were inoculated per each isolate × cultivar combination, i.e. 200 fruits per cultivar, including the controls. After inoculation, fruits were incubated for 7 days in humid chamber at 23±1 °C, with a photoperiod of 16 h of light and 8 h of dark and 80% relative humidity. Symptom severity on inoculated fruits was rated daily using an empirical scale from 0 to 6 (Talhinhas et al., 2015) and the relative area under disease progress curve (rAUDPC) was calculated according to Riolo et al. (Riolo et al., 2022). The experiment was repeated three times. Results of the three experiments were pooled together as they did not differ significantly from each other.

### 2.3 Secondary metabolites analysis by HPLC-QTOF

To extract the secondary metabolites produced by *Colletotrichum* species on olives during the infection process, flesh fragments (5 mm diameter) were sampled from inoculated olives at three different time intervals, i.e. 1, 3 and 7 days post inoculation (dpi). Fragments were excised from around the inoculation site and transferred into 15 ml Eppendorf tubes (three fragments from three distinct olives of the same batch per tube). For each batch, three replicates (tubes) were processed at each sampling time. Pulp fragments were weighted and a volume of methanol (MeOH) was added in a 1:5 (w/v) ratio. The tubes were incubated at room temperature and continuous stirring (150 rpm) for 24 hours. After incubation, the extract was centrifugated at 4,000 rpm for 15 min at 4°C, the supernatant was filtered through a 0.22 μm filter and injected for HPLC–QTOF analysis. All the analyses were performed in triplicate. The extraction of metabolites produced by *Colletotrichum* species on agar media (PDA and OA) was performed with the same method with a few modifications. Fungal isolates were grown in Petri dishes at 23 ±1 °C for 7 days. Then dishes were transferred into sterile urine beakers and methanol was added in a 1:2 (w/v) ratio. Then the samples were incubated at room temperature and stirring (150 rpm) for 24 hours. After incubation, the extract was centrifugated at 4000 rpm for 15 min at 4°C, the supernatant was filtered through a 0.22 μm filter and injected for HPLC–QTOF analysis. All the analyses were performed in triplicate. An HPLC Agilent 1200 (Agilent Technologies, Santa Clara, CA, USA) was used for the chromatographic analysis, it consisted of an automatic sampler, a binary pump and a vacuum degasser. Injection volume was 5 μL and the analysis was performed in 25 minutes. The separation of the analytes was performed using a Gemini C18 column (50 mm × 2 mm, 110 Å and particle size 3 μm) (Phenomenex, Palo Alto, CA, USA). Mobile phases consisted of water (solvent A) and acetonitrile (solvent B), both with 0.1% formic acid. The elution flow rate was 0.3 mL/min and the elution gradient was as follows: 0 min, 5% B; 30 min, 95% B and 35 min, 5% B. Mass spectrometry analyses were performed using a QTOF (6540 Agilent Ultra High-Definition Accurate Mass, Agilent Technologies, Santa Clara, CA, USA), coupled to an Agilent Dual Jet Stream electrospray ionization (Dual AJS ESI) interface operating in positive ion mode. Optimized mass spectrometry parameters included: capillary voltage 3.5 kV; fragment voltage 175 V; drying gas flow (N2) 8 L/min, temperature 350 °C; collision energy 10, 20 and 40 eV, nebulizer pressure 30 psi. Data analysis was performed by MassHunter Qualitative Analysis Software B.08.00 (Agilent Technologies, Santa Clara, CA, USA).

### 2.4 Statistical analysis

To compare the severity of symptoms on olives of different cultivars all the data were normalized by square root transformation and then subjected to analysis of variance (ANOVA), followed by Tukey’s honestly significant difference (HSD) test as a post hoc test (R software). The analytical data obtained by HPLC-QTOF were log10 transformed before statistical analysis. The secondary metabolites obtained from olive cultivar x *Colletotrichum* species combinations at each sampling time (1, 3 and 7 dpi) were compared using the LSD test at P ≤ 0.05, after analysis of variance (one-way ANOVA). Relationships among olive cultivar x *Colletotrichum* species combinations were analyzed using Pearson’ s correlation coefficient analysis. All the above statistical analyses were performed using RStudio v.1.2.5 (R). MetaboAnalyst 5.0 software (Pang et al., 2021) was used for principal component analysis (PCA) using log10 transformed data. The features included were log transformed and mean centred.

## 3. Results

In artificial inoculation assays *Colletotrichum acutatum* was the most virulent and *C. karsti* the least virulent of the four *Colletotrichum* species tested, as expected (Figure 1).

**Figure 1.**
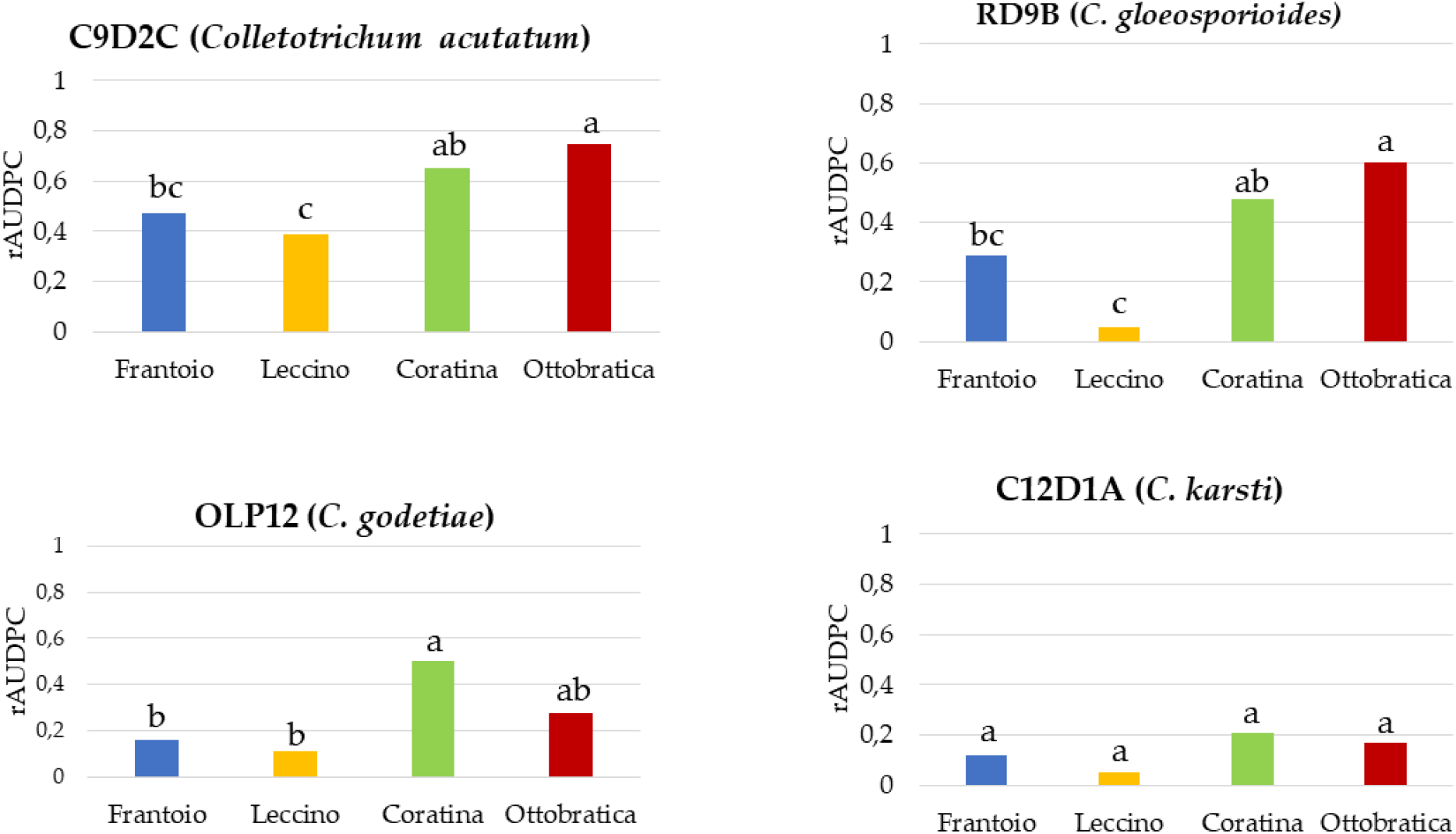
Mean olive anthracnose intensity over time (relative area under the disease progress curve [rAUDPC]) on drupes of nine olive cultivars inoculated with isolates of *Colletotrichum acutatum*, *C. gloeosporioides*, *C. godetiae* or *C. karsti* (one isolate per species). Severity of symptoms on inoculated olive drupes was assessed daily for 7 days and rated on a 0 to 6 scale according to the size of necrotic lesion and the abundance of fungal sporulation. Values sharing a common letter are not statistically different according to Tukey’s honestly significant difference (HSD) test (*p* ≤ 0.05).

Moreover ‘Ottobratica’ and ‘Coratina’ were confirmed to be very susceptible to the infection of *Colletotrichum* species, while both ‘Leccino’ and ‘Frantoio’ were once more confirmed to be relatively resistant also to the most aggressive *Colletotrichum* species. No significant difference in susceptibility was noticed among the four olive cultivars inoculated with *C. karsti* as this *Colletotrichum* species was poorly aggressive on all tested cultivars.

Overall, the analysis by HPLC-QTOF revealed 45 secondary metabolites on both fruits of different olive cultivars inoculated singularly with four *Colletotrichum* species and axenic cultures of *Colletotrichum* isolates on agar media (Table 2).

**Table 2.**
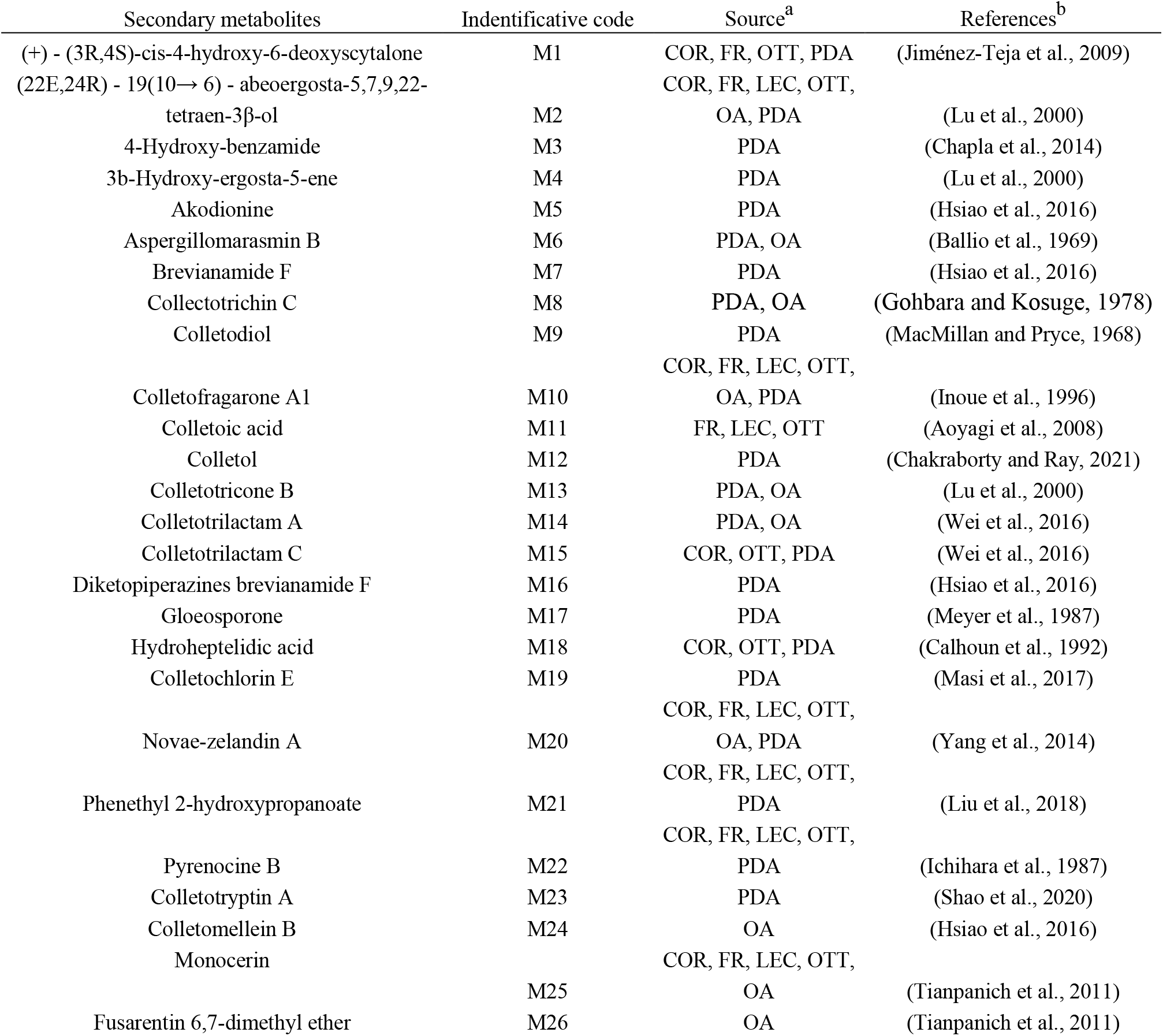
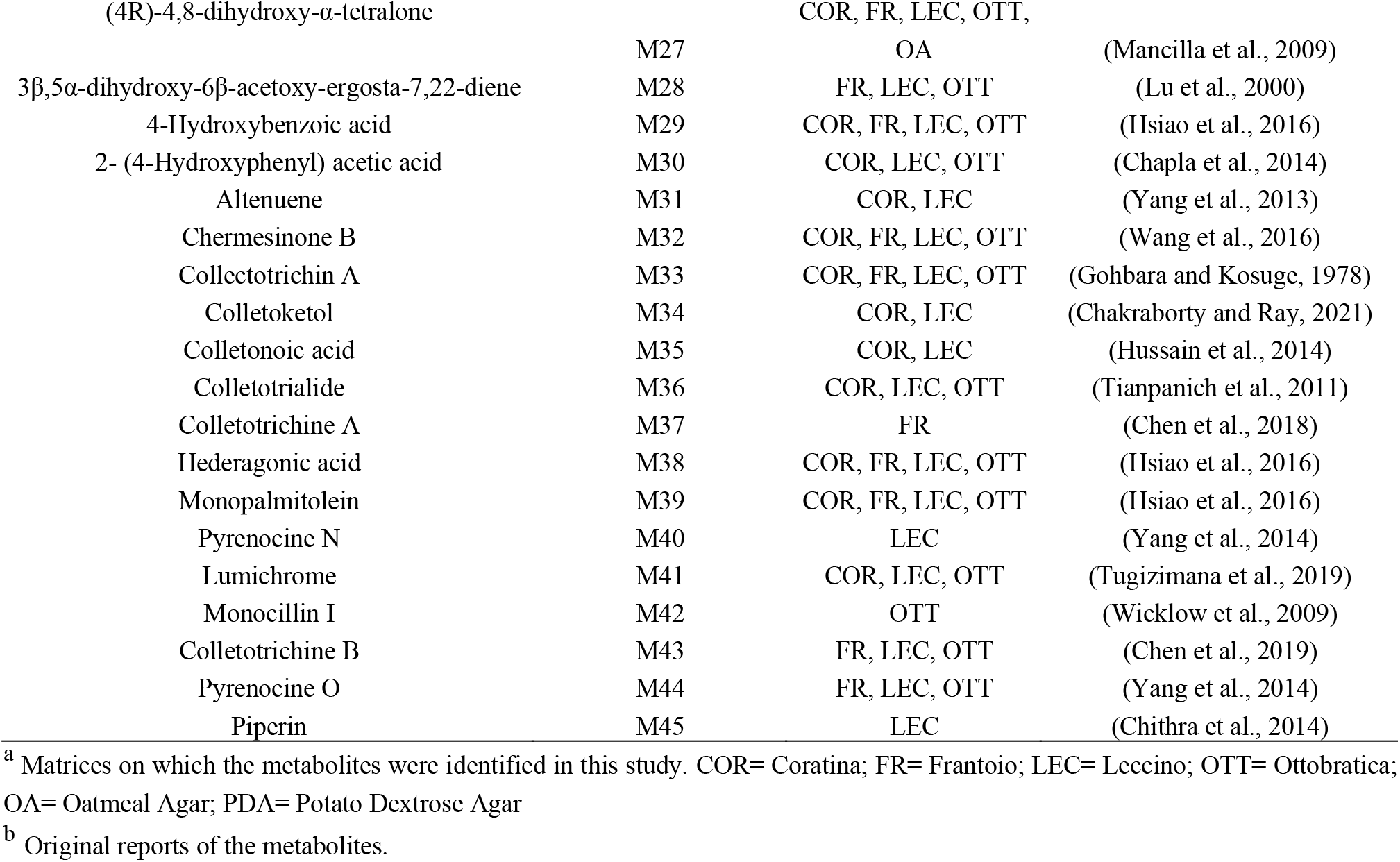
Secondary metabolites of *Colletotrichum* identified in this study.

Not all secondary metabolites produced on olives were detected in vitro and viceversa. Overall 29 metabolites were detected in extracts from infected olives, 26 in extracts from pure cultures of *Colletotrichum* species on agar media and of these only 10 were found in extracts from both inoculated olives and pure cultures while 19 were detected exclusively on olives. Only 16 metabolites were found exclusively in pure cultures on artificial media; of these four were detected in both OA and PDA extracts, 10 exclusively in PDA extracts and two exclusively in OA extracts. The identification of the metabolites was supported by a purpose-built database, based on the secondary metabolites produced by *Colletotrichum* species reported in the literature (Table 2). The absence of these metabolites in extracts from control non-inoculated olive fruits confirmed the 45 identified metabolites were of fungal origin (Figure 2 and Figure 3).

**Figure 2.**
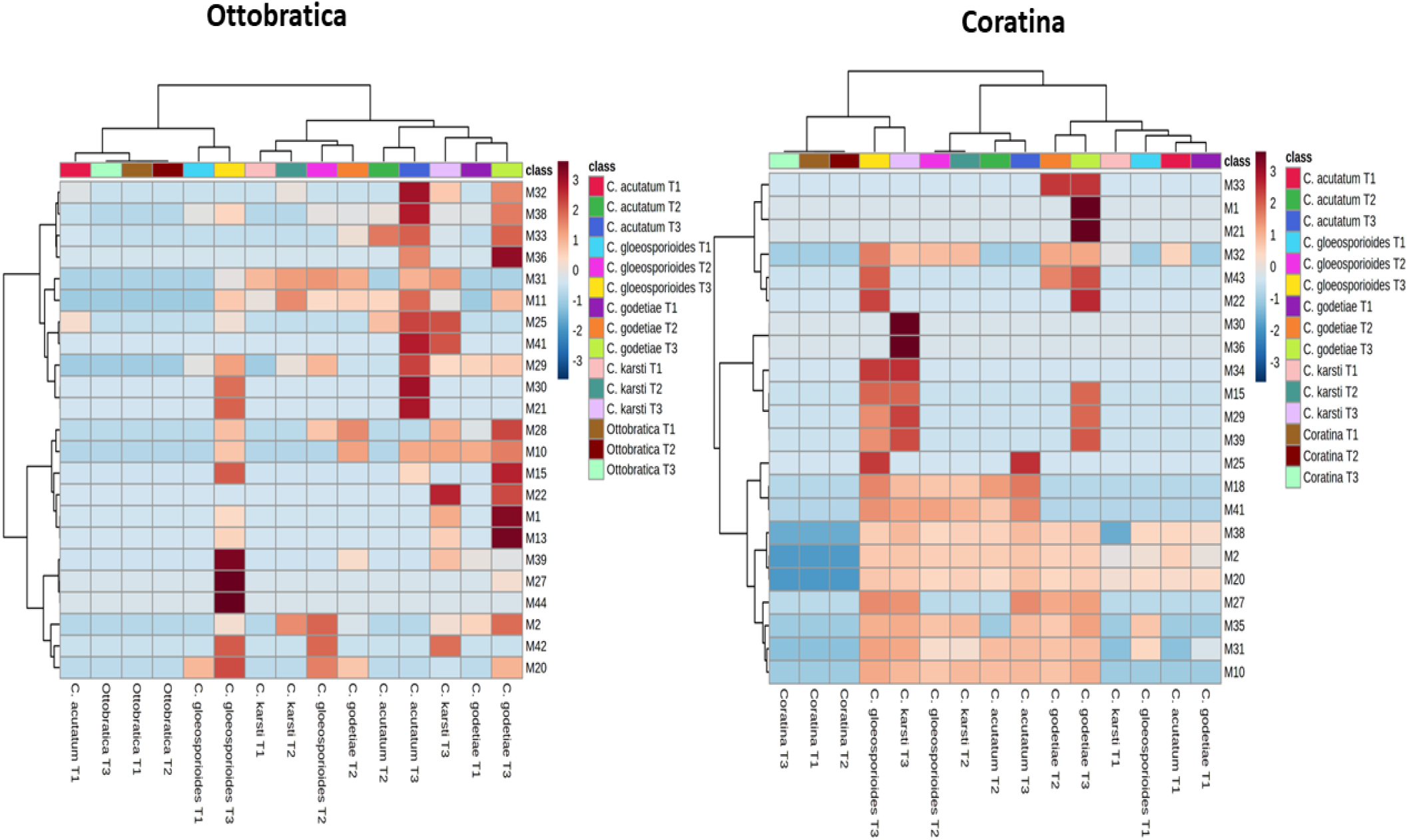
Heat map of metabolites produced by *Colletotrichum* species on olives of ‘Ottobratica’ and ‘Coratina’. Colors indicate the relative abundance (logarithmic scale) of metabolites produced in each olive cultivar x *Colletotrichu*m species combination during the infection process, at different time intervals after inoculation (T1, T2 and T3, corresponding to one, three and seven days post inoculation, respectively).

**Figure 3.**
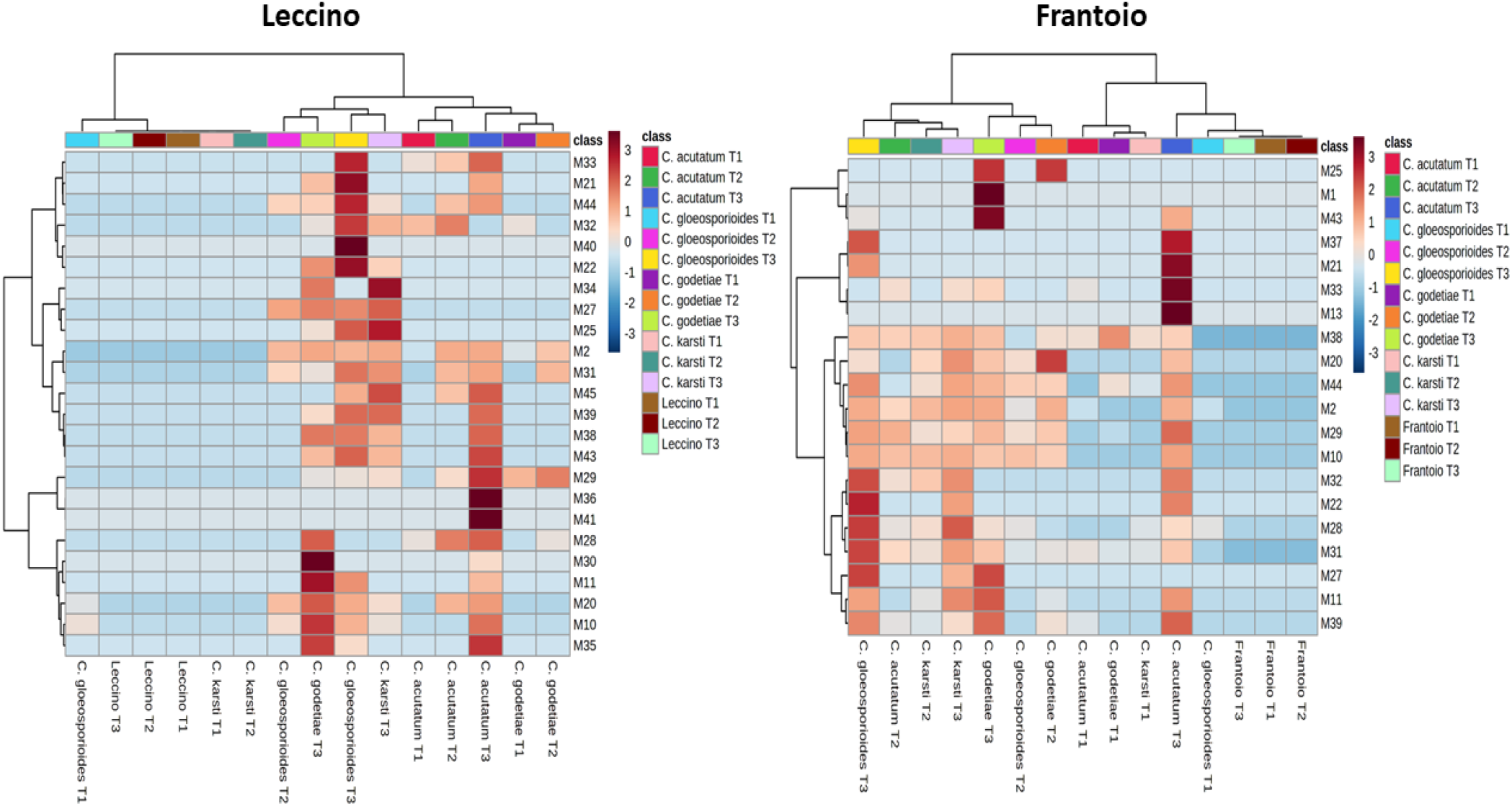
Heat map of metabolites produced by *Colletotrichum* species on olives of ‘Leccino’ and ‘Frantoio’. Colors indicate the relative abundance (logarithmic scale) of metabolites produced in each olive cultivar x *Colletotrichu*m species combination during the infection process, at different time intervals after inoculation (T1, T2 and T3, corresponding to one, three and seven days post inoculation, respectively). Brown and blue represent high and low abundance, respectively. Non inoculated olives of each cultivar at T1, T2 and T3 were included as controls.

The semi-quantitative heat map method made it possible to compare the spectra of secondary metabolites produced by the *Colletotrichum* species on each olive cultivar at three different time intervals after inoculation. In general, the production of secondary metabolites in all olive cultivars inoculated singularly with each of the four *Colletotrichum* species peaked at 7 dpi while the metabolite profiles of non-inoculated controls, i.e. olives treated only with water, at three time intervals after wounding, clustered together, as no detectable amounts of fungal metabolites were recorded on these olives (Figure 2 and Figure 3). At T1 (1 dpi), the metabolite profiles of ‘Ottobratica’, ‘Coratina’ and ‘Frantoio’ olives inoculated with *C. acutatum*, *C. gloeosporioides*, *C. godetiae* or *C. karsti*, showed traces of one or few of the fungal metabolites found in the respective metabolite profiles at T2 (3 dpi) and T3 (7dpi). Conversely, the metabolite profiles of ‘Leccino’ olives inoculated with *C. karsti*, at both T1 and T2 did not differ from the non-inoculated controls as no amount of fungal metabolites was detected, while the profile of olives of this olive cultivar inoculated with *C. acutatum*, *C. gloeosporioides* or *C. godetiae*, at T1 showed only traces of one (*C. gloeosporioides* and *C. godetiae*) up to a maximum of three (*C. acutatum*) of the fungal metabolites found in the respective metabolite profiles at T2 and T3. In detail, at T3 (7 dpi) in olives of ‘Ottobratica’ inoculated with *C. acutatum* and *C. godetiae* the most abundant secondary metabolites were M32, M38, M33, M36 and M10. (Figure 2). Monocerin (M25), together with Lumichrome (M41) were recorded on drupes inoculated with *C. acutatum* or *C. karsti*. High amounts of simple phenolic compounds, such as 4-hydroxybenzoic acid (M29) and 2-(4-Hydroxyphenyl) acetic acid (M30), were detected on olives of this cultivar inoculated with *C. acutatum* or *C. godetiae*. At T3, particularly high amounts of M28, M10, M15, M22, M1 and M13 metabolites were detected in olives inoculated with *C. godetiae*, while M27, M39 and M44 were the prevalent fungal metabolites in olives inoculated with *C. gloeosporioides*. At T3 the metabolite M10 (Colletofragarone A1) was detected on all olives inoculated with one of the four *Colletotrichum* species. On olives inoculated with *C. godetiae* this last metabolite was already detected at T1 (1 dpi). As for ‘Coratina’ (Figure 2), the metabolites M2, M10, M20, M27, M31, M35 and M38 were detected in all the metabolite profiles of olives inoculated with *C. acutatum*, *C. gloeosporioides*, *C. godetiae* or *C. karsti* at T3 (7 dpi) as well as in the metabolite profile of olives inoculated with *C. godetiae*, at T2 (3dpi). All these metabolites, with the only exception of M27, were also detected at T2 in the profile of olives inoculated with *G. gloeosporioides* and *C. karsti*, and all, except M27 and 35, were present in the profile of olives inoculated with *C. acutatum*, at T2. In the metabolite profile of olives inoculated with *C. godetiae* at T3, two fungal metabolites, M1 [(+) - (3R,4S)-cis-4-hydroxy-6-deoxyscytalone)] and M21 (Phenethyl 2-hydroxypropanoate), stood out for their high amount. In the profile of these olives also the metabolites M22 (Pyrenocine B), M33 (Collectotrichin A) and M43 (Colletotrichine B) were highly represented. Collectotrichin A was extracted in a relatively high amount from olives inoculated with *C. godetiae* even at T2 (3 dpi). The metabolites M22 and M43 were also detected in a relative high amount in the profile of olives inoculated with *C. gloeosporioides*, at T3. The metabolites M30 and M36 were the most abundant in the metabolite profile of olives inoculated with *C. karsti*. In the profile of these olives also the metabolites M15, M29, M34 and M39 were detected in a relative high amount. At T3, the last four metabolites and the metabolite M25 were also abundant in the profile of olives inoculated with *C. gloeosporioides*. The most abundant fungal metabolite in the profile of olives inoculated with *C. acutatum* at T3 was M25 (Monocerin), which at T3 was also present in a relatively high amount in the metabolite profile of olives inoculated with *C. gloeosporioides*. Overall, the highest number of fungal metabolites was detected in the profile of olives inoculated with *Colletotrichum gloeosporioides* and *C. godetiae*.

As for ‘Leccino’, the most resistant cultivar tested, at T3 (7 dpi) the metabolite profiles of olive batches inoculated with the four *Colletotrichum* species were substantially different as regards the relative abundance of each fungal metabolite (Figure 3). The highest number of metabolites (in total 18 per profile) was found in the profile of olives inoculated with *C. acutatum*, *C. gloeosporioides* and *C. godetiae*, while the profile of olives inoculated with *C. karsti* consisted of only 15 metabolites. The metabolite profile of olives inoculated with *C. acutatum* was characterized by a high amount of M36 (Colletotrialide) and M41 (Lumichrome). Also high amounts of metabolites 29, 35, 43 and 45 were detected. The metabolite profile of olives inoculated with *C. gloeosporioides* was characterized by a high amount of M40 (Pyrenocine N). Other metabolites of this profile present in a relatively high amount included M21, M22, M32, M33 and M44. The metabolite profile of olives inoculated with *C. godetiae* was characterized by a high amount of M30 [2-(4-Hydroxyphenyl) acetic acid]. Other metabolites of this profile present in a relatively high amount included M10, M11 and M35. Finally, the most abundant fungal metabolites in the profile of extracts from olives inoculated with *C. karsti* were M25, M27, M34 and M45.

Also for ‘Frantoio’, a resistant cultivar like ‘Leccino’, the metabolite profiles of olives inoculated with the four *Colletotrichum* species at three diverse time intervals after inoculation (T1, T2 and T3, corresponding to 1, 3 and 7 dpi) can be separated into three distinct groups (Figure 3). The first group comprised the profiles of non-inoculated controls at T1, T2 and T3, which did not show the presence of any fungal secondary metabolite. The second group comprised the profile of olive batches inoculated with *C. acutatum*, *C. gloeosporioides*, *C. godetiae* or *C. karsti* at T1, which showed traces of only one or a maximum of four fungal secondary metabolites. The third group comprised the profiles of all other olive batches. Also for this olive cultivar the highest amount of fungal metabolites was detected in olives inoculated with the four *Colletotrichum* species processed 7 dpi (T3). The richest in fungal metabolites (17 in total) was the profile obtained from olives inoculated with *C. acutatum*, at T3. This profile was characterized mostly by high levels of M13 (Colletotrichone B), M21 (Phenethyl 2-hydroxypropanoate), M33 (Colletotrichin A) and M37 (Colletotrichine A). Relatively high amounts of M22, M29, M30, M32 and M40 were also present in this profile. The profile obtained from olives inoculated with *C. godetiae* comprised 15 diverse fungal metabolites and was characterized by high levels of M1 [(+) - (3R,4S)-cis-4-hydroxy-6-deoxyscytalone] and M43 (Colletotrichine B). Relatively high amounts of M11, M27 and M39 were also present in this profile. Similar to the profile associated to the infection by *C. godetiae*, the metabolite profile from olives inoculated with *C. gloeosporioides* at T3 comprised in total 15 metabolites and only traces of a sixteenth metabolite (M43). The most abundant metabolite of this profile was M22 (Pyrenocyne B). Other metabolites detected in a relatively high amount included M27, M28, M31 and M32. The most abundant fungal metabolite in the profile of olives ‘Frantoio’ inoculated with *C. karsti*, at T3 was M28 (3β,5α-dihydroxy-6β-acetoxy-ergosta-7,22-diene). The metabolite profile of olives inoculated with *C. karsti* comprised in total 14 diverse metabolites and shared with the profile from olives inoculated with *C. gloeosporioides* 13 metabolites, including M2, M10, M11, M20, M22, M27, M28, M29, M31, M32, M38, M39 and M44.

In order to evaluate the production of secondary metabolites by various *Colletotrichum* species on different olive cultivars, a PCA analysis of data was performed, based on the secondary metabolites identified (Table 2) on olive cultivars inoculated with the four diverse *Colletotrichum* species, at three distinct time intervals after inoculation (T1, T2 and T3).

Figure 4 shows the clustering of the inoculated olives at different time intervals (score plot) and the metabolite trends (loading plot) in different quadrants (I, II, III, IV); the sum of the principal component values reached 61.8% of the total variance. PC1 represented 44.5% and PC 2 17.3% of the total variance.

**Figure 4.**
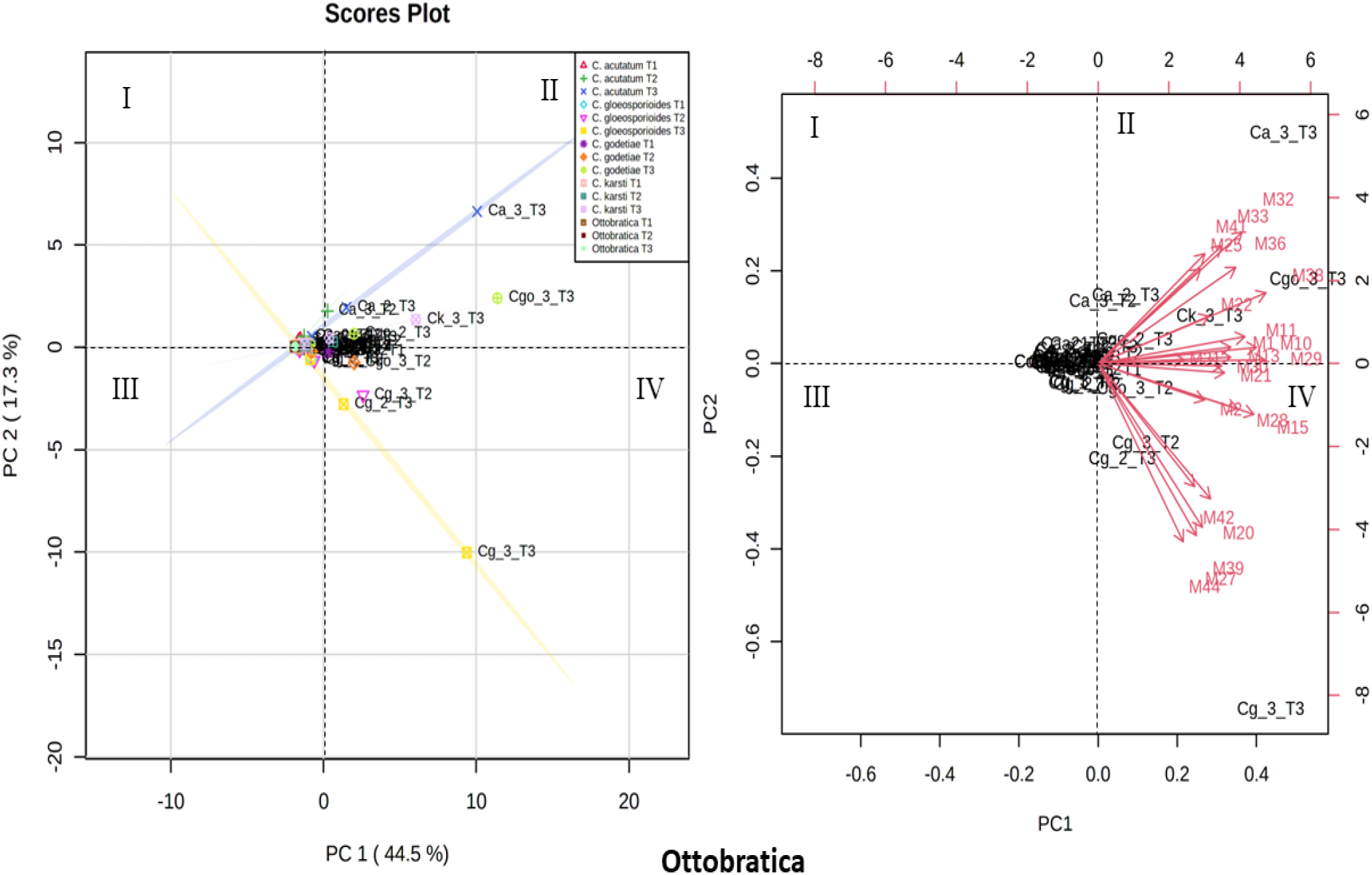
Principal Component Analysis (PCA), scores plot (a) and loading plot (b), based on secondary metabolites produced by *C. acutatum* (Ca), *C. gloeosporioides* (Cg), *C. godetiae* (Cg) and *C. karsti* (Ck) on the cultivar Ottobratica at different time intervals after inoculation (T1, T2 and T3).

In the score plot of the cultivar Ottobratica (Figure 4), the value 0 was recorded for all samples at both T1 (1 dpi) and T2 (3 dpi), while each group was split at T3 (7 dpi), based on the secondary metabolites produced. Samples infected by *Colletotrichum acutatum* (Ca_1,2,3_T3), *C. karsti* (Ck_1,2,3_T3) and *C. godetiae* (Cgo_1,2,3_T3) clustered in the quadrant II. Samples infected by *C. gloeosporioides* (Cg_1,2,3_T3), with the only exception of replicate 1, tend towards quadrant IV. In the loading plot, metabolites M44, M39, M20 and M42 tend towards quadrant IV, clustering the *C. gloeosporioides* samples, at T3 (7 dpi). By contrast, metabolites M32, M33, M41, M25, M36, M38, M22, M11, M19, M1and M29 cluster the samples infected by *C. acutatum*, *C. godetiae* and *C. karsti* at T3 (Figure 4). As for the cultivar Coratina, the sum of the principal component values reached 67% of the total variance. PC1 represented 48.4% and PC 2 18.6% of the total variance. On the score plot, drupes inoculated with *C. godetiae* at T2 and T3 were grouped in the II quadrant. By contrast, olives inoculated with *C. karsti*, *C. gloeosporioides* and *C. acutatum* at T3. clustered in the IV quadrant. The rest of the samples tended towards the centre. The loading plot indicates that the secondary metabolites M21, M35, M10, M38, M34, M30, M18 and M41 clustered the samples at infected by *C. karsti*, *C. acutatum* and *C. gloeosporioides*, at T3. The rest of the samples cluster around the value 0 (Figure 5).

**Figure 5.**
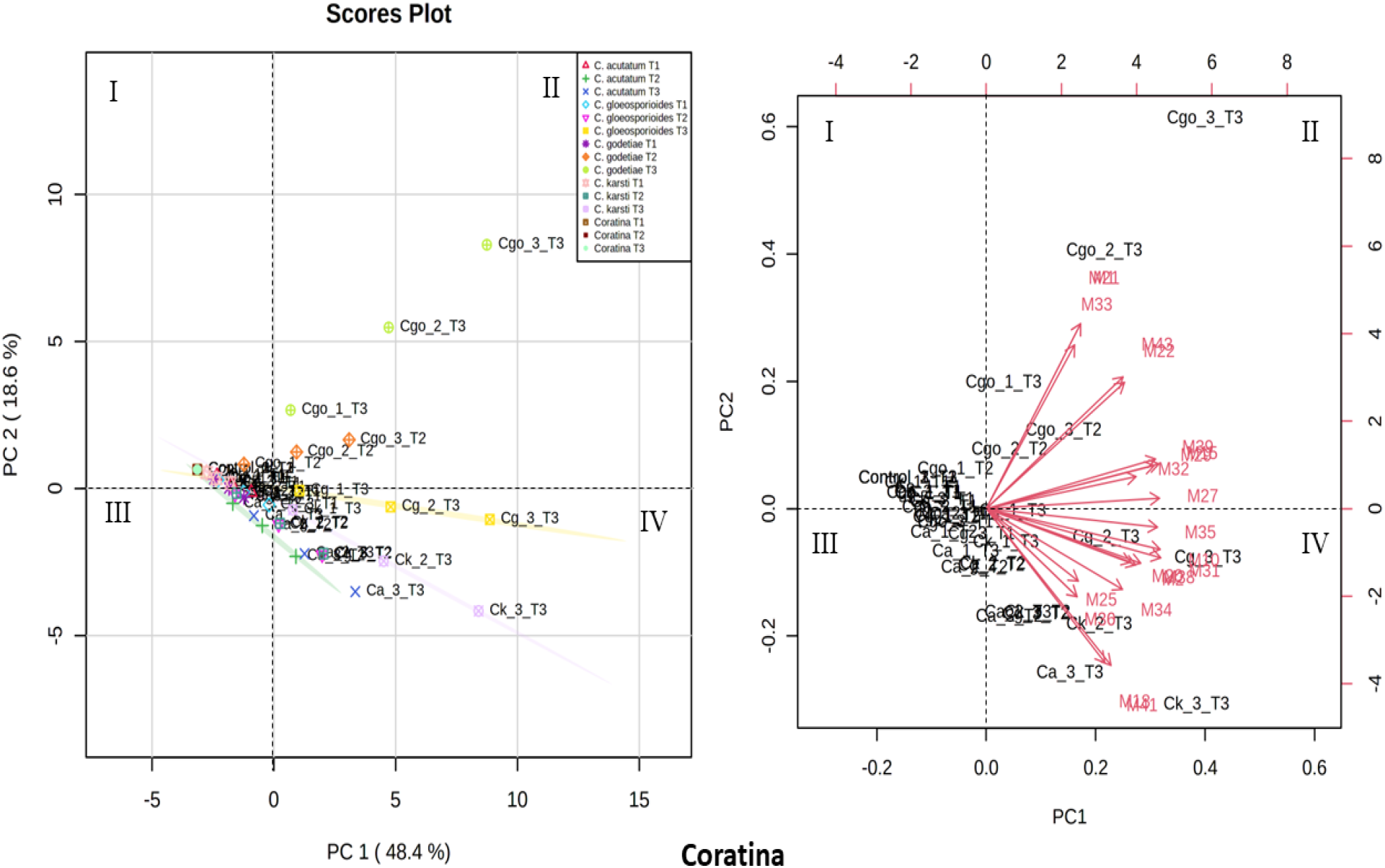
Principal Component Analysis (PCA), scores plot (a) and loading plot (b), based on secondary metabolites produced by *C. acutatum* (Ca), *C. gloeosporioides* (Cg), *C. godetiae* (Cg) and *C. karsti* (Ck) on the cultivar Coratina at different time interval after inoculation (T1, T2 and T3).

The PCA of the Frantoio cultivar recorded a sum of components equal to 71.4% of the total variance, of which PC1 represented 54.2% and PC2 17.2%. Analysing the score plot in Figure 6, samples inoculated with *C. godetiae* at T2 and T3 cluster within quadrant II, while olives inoculated with *C. gloeosporioides* and those inoculated with *C. acutatum*, at T3 cluster in quadrant IV. The rest of the samples, at T1 and T2, tend to cluster around the 0 point. As regards the direction of the secondary metabolites in the loading plot, M11 and M25 tend to cluster the samples infected by *C. godetiae* (C.go_1,2,3_T2,T3) at T2 and T3 towards quadrant II. By contrast, secondary metabolites M38, M2, M11, M44, M39, M10, M31 and M29 cluster samples infected by *C. karsti* at T3 in quadrant II. Morover, secondary metabolites M28, M33, M22, M32, M13, M21 and M37 tend towards quadrant IV, clustering the samples inoculated with *C. acutatum* (Ca_1,2,3_T3) and *C. gloeosporioides* (Cg_1,2,3_T3) at T3 (Figure 6).

**Figure 6.**
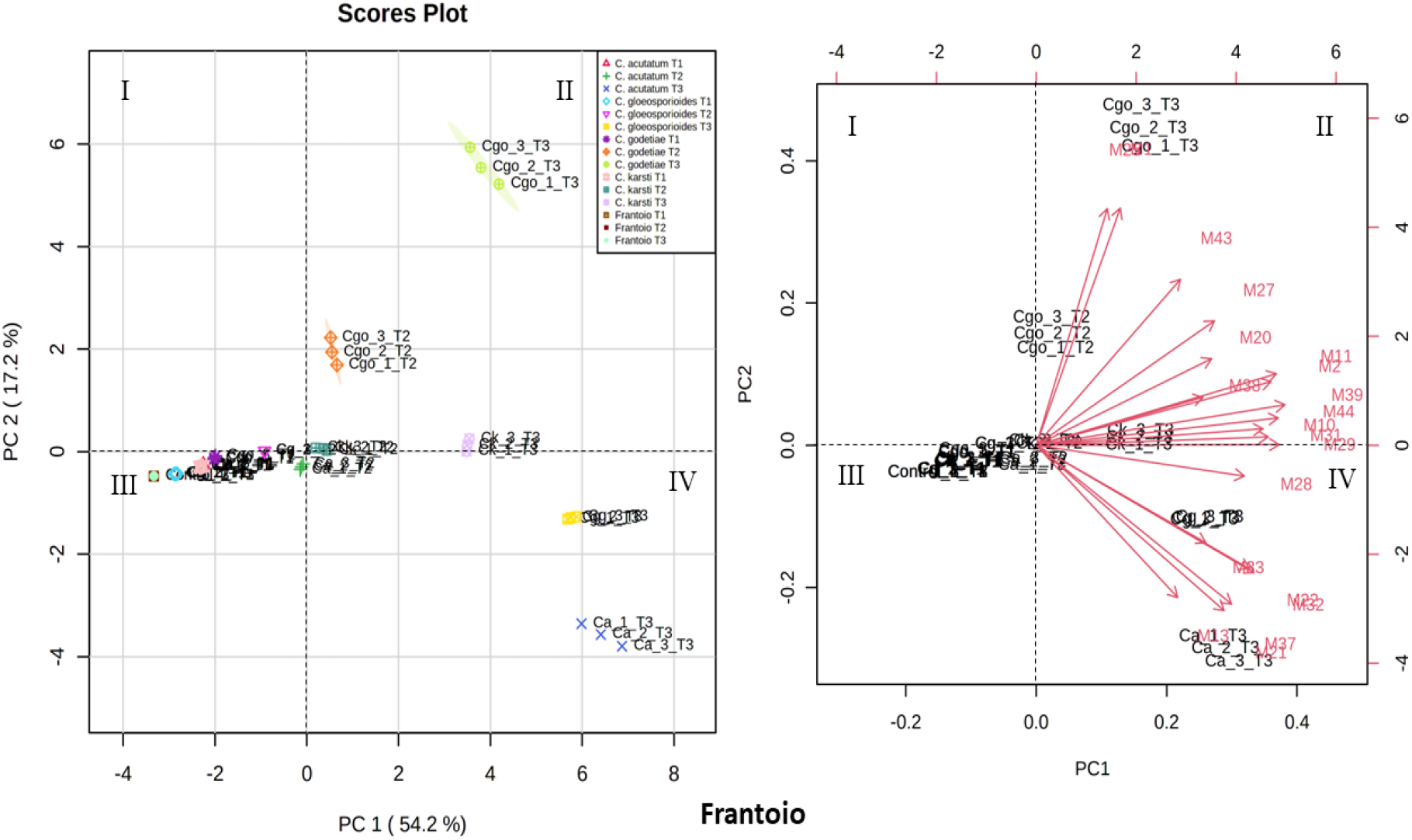
Principal Component Analysis (PCA), scores plot (a) and loading plot (b), based on secondary metabolites produced by *C. acutatum* (Ca), *C. gloeosporioides* (Cg), *C. godetiae* (Cg) and *C. karsti* (Ck) on olives of the cultivar Frantoio at different time intervals after inoculation (T1, T2 and T3).

Finally, as regards the cultivar Leccino, it presented a sum of components equal to 69.1% of the total variance, with PC1 and PC2 percentage values of 52.2 and 16.9, respectively (Figure 7). Analysing the quadrants of the score plot, samples inoculated with *C. acutatum* and *C. godetiae* at T3 clustered in quadrant II, differently from olives inoculated with *C. karsti* and *C. godetiae* at T3, which grouped in quadrant IV. Conversely, olives inoculated with *C. godetiae* at T2 clustered in quadrant I. Olives inoculated with *C. gloeosporioides* at T2 and *C. acutatum* at T1 grouped in quadrant III. Loading plot analysis revealed that the secondary metabolites M36, M41, M28, M29, M35, M30, M10, M11, M20, M43 and M38 tended towards quadrant II, grouping the drupes inoculated with *C. acutatum* and *C. godetiae* at T3, whereas olives inoculated with *C. karsti* and *C. gloeosporioides* at T3 grouped on the basis of secondary metabolites M38, M45, M39, M31, M44, M21, M34, M27, M22, M32, M40 and M25.

**Figure 7.**
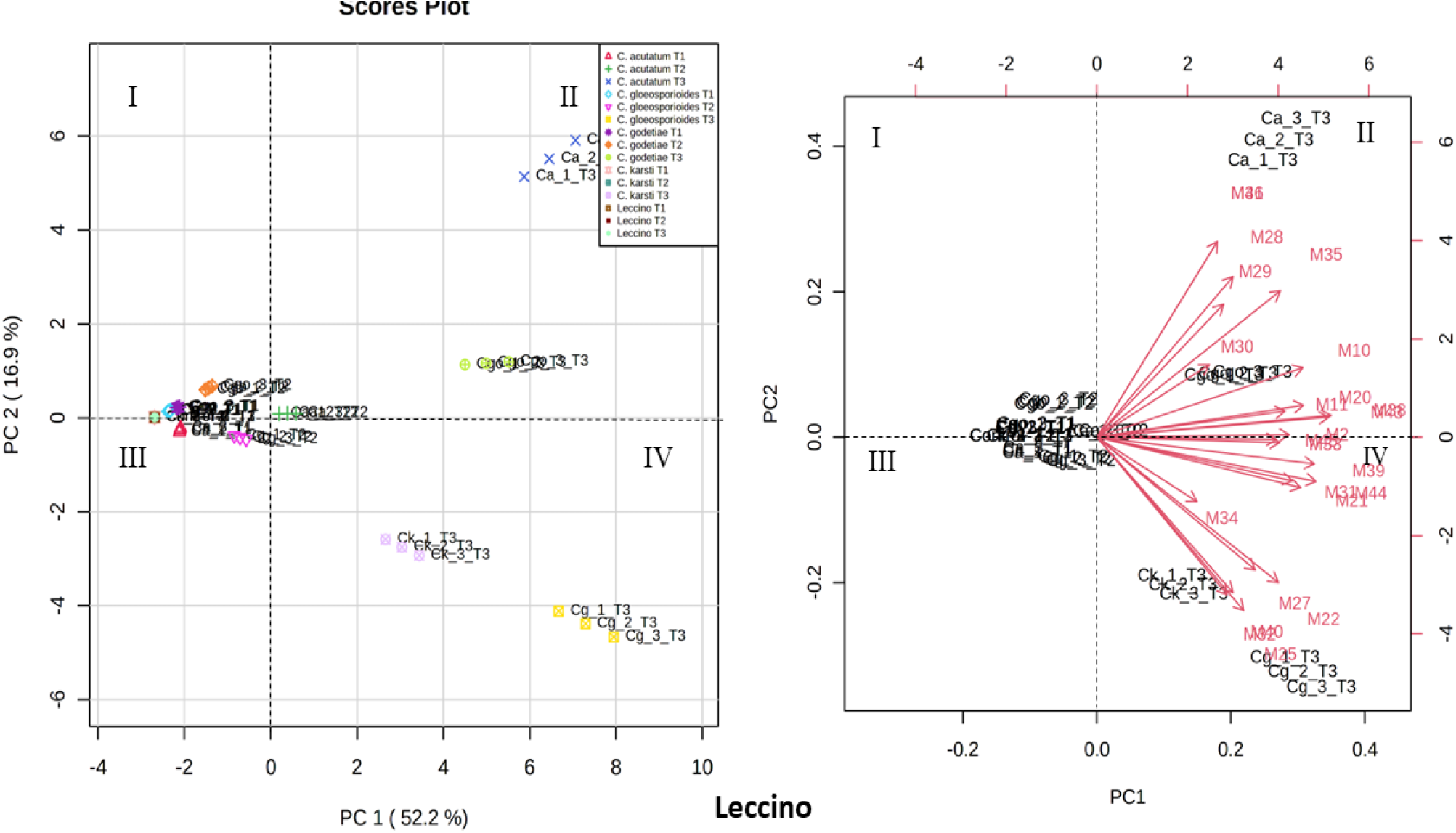
Principal Component Analysis (PCA), scores plot (a) and loading plot (b), based on secondary metabolites produced by *C. acutatum* (Ca), *C. gloeosporioides* (Cg), *C. godetiae* (Cg) and *C. karsti* (Ck) on olives of the cultivar Leccino at different time intervals after inoculation (T1, T2 and T3).

The metabolites produced by *C. acutatum*, *C. gloeosporioides*, *C. godetiae* and *C. karsti* on olives of different cultivars were compared with those produced by these four species of *Colletorichum* on agar media (PDA and OA). On PDA, heat map analysis revealed abundant production of the metabolites M4 and M8 by *C. karsti* isolates (CAM and C12D1A, respectively). In contrast, the secondary metabolites M3 and M12 were only produced by *C. karsti* isolate C12D1A sourced from olive drupes. Medium to high values of the secondary metabolites M19, M5, M13 and M14 were recorded from *C. godetiae* isolates OLP10 and OLP12. M14 was also detected in medium to high quantities in the extracts of the *C. karsti* isolate C12D1A. The metabolites M5, M13, M17 and especially M1 were produced from the *C. acutatum* isolate UWS149. Both *C. acutatum* isolates produced in abundance the metabolites M23, M18, M9, M15 and M20. In contrast, the *C. gloeosporioides* isolates were characterized by the production of M2, M6, M21, M22, M17, M20 and M19. On Oatmeal agar (OA), a lower number of metabolites were produced than on PDA and some of these metabolites were different from those on PDA. *Colletotrichum karsti*, for example, produced M13 on OA but not on PDA. Conversely, both isolates of *C. gloeosporioides* produced M20 on PDA but not on OA. This secondary metabolite was also produced by isolates of *C. karsti* on OA (Figure 8).

**Figure 8.**
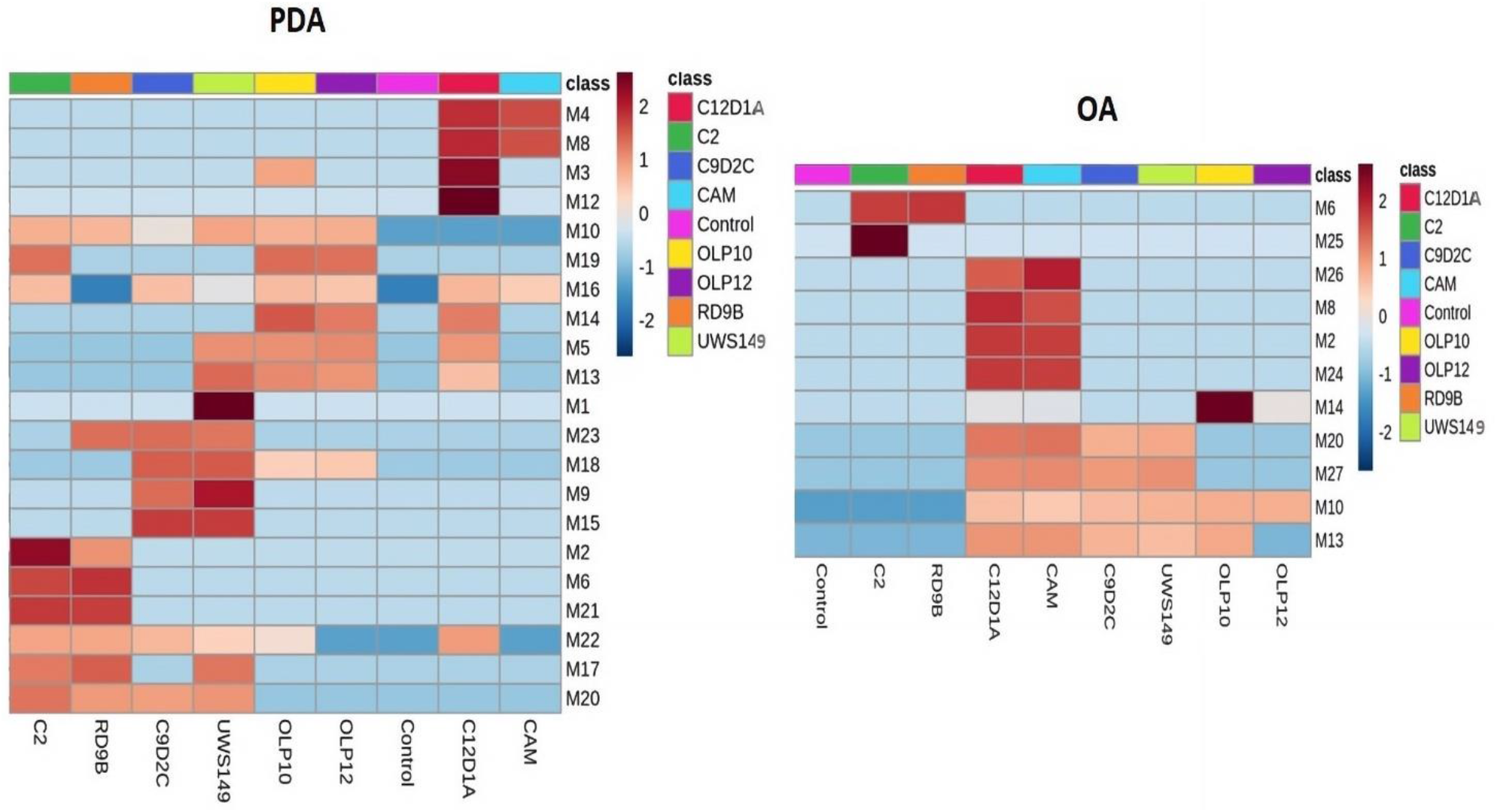
Heat map of metabolites produced by *Colletotrichum* species in axenic cultures on two diverse agar media (PDA and OA). Colors indicate the relative abundance (logarithmic scale) of metabolites produced in each medium. Brown and blue represent high and low abundance, respectively.

In detail the only metabolites produced by *C. gloeosporioides* on OA were M6 and M25. By contrast, both species of *C. karsti* on OA produced abundantly the metabolites M26, M8, M2 and M24, *Colletotrichum acutatum* producesd more metabolites on PDA, while *C. karsti* produced more metabolites on OA (Figure 8).

## 4. Discussion

This study is the first to characterize the secondary metabolites released by diverse *Colletotrichum* species associated with OA both *in vitro* and during the infection process of olive fruits. It showed that these fungal pathogens produce a wide array of secondary metabolites both *in vitro* and *in planta*. In total, 45 diverse metabolites were identified; of these 29 were detected on infected fruits, 26 in axenic cultures and 10 on both fruits and axenic cultures. Overall, 10 out of 26 secondary metabolites produced *in vitro* were found exclusively in axenic cultures grown on PDA, while two were found exclusively in axenic cultures on OA and four on both culture media, indicating the metabolite profile of *Colletortrichum* species was strongly influenced by the matrice. Each *Colletotrichum* species produced a different spectrum of metabolites. Moreover the metabolite profile of each species varied both qualitatively and quantitatively depending of the type of matrices. Substantial differences in the metabolic profile of the same *Colletotrichum* species were observed in extracts from both different artificial culture media and olive cultivars. The secondary metabolites identified on olive fruits inoculated with *Colletotrichum* included compounds with phytotoxic and/or cytotoxic activity such as Chermesinone B and Colletonoic acid, produced in a relatively high amounts in olives infected by *C. acutatum* and *C. godetiae* and possessing antibacterial, antifungal and antialgal properties (Hussain et al., 2014; Wang et al., 2016). Other secondary metabolites with phytotoxic and/or cytotoxic activity identified in this study, included Colletofragarone A1 and Pyrenocine B, also among the secondary metabolites produced by *C. godetiae* on infected olives, which according to the literature are phytotoxic (Kim and Shim, 2019; Yang et al., 2014), Pyrenocine O, produced by *C. gloeosporioides* on olives of ‘Ottobratica’ and belonging to a class of compounds with several biological properties, including phytotoxicity, cytotoxicity and antifungal efficacy (Yang et al., 2014), Pyrenocine B and Colletotrichine B, produced in abundance by both *C. gloeosporioides* and *C. godetiae* on olives of ‘Coratina’ and known for their phytotoxicity (Chen et al., 2019; Yang et al., 2014), the polyketide metabolite Monocerin produced by *C. godetiae* on olives of ‘Leccino’ and reported originally as a phytotoxic metabolite produced by the fungus *Exserohilum turcicum* (formerly known as *Helminthosporium turcicum*), the causal agent of northern corn leaf blight disease in maize (Cuq et al., 1993). This corroborates the hypothesis of an involvement of secondary metabolites of *Colletotrichum* species in the pathogenesis of OA. Furthermore, if it were demonstrated that biologically active fungal secondary metabolites produced on infected olives would contaminate the oil during the extraction process this might have toxicological implications. Examining the variability of secondary metabolite profiles resulting from the interaction *Colletotrichum* species x olive cultivar and comparing the differences between susceptible and resistant cultivars or between aggressive and less virulent *Colletorichum* species, no obvious difference in the ability of producing specific metabolites can be correlated with the symptom severity. By contrast, substantial differences were observed in the dynamics of metabolite profiles in resistant (‘Leccino’ and ‘Frantoio’) and susceptible (‘Ottobratica’ and ‘Coratina’) olive cultivars, irrespective of the *Colletotrichum* species involved. Not surprisingly, the amount and the number of metabolites produced in infected olives increased dramatically over the course of infection but this process has been considerably slowed down in resistant olive cultivars. These trends were confirmed my multicomponent analysis. In the light of the results of this study showing that several different bioactive secondary metabolites are produced on olives infected by *Colletotrichum* species the assumption that Aspergillomarasmin produced in rotten fruits is responsible for the symptoms on twigs and leaves (Moral et al., 2021) appears as an oversimplification. As a matter of fact, Aspergillomarasmin, first identified by Ballio et al. (Ballio et al., 1969) in culture fluids of *C. gloeosporioides s.l*., in this study was extracted (in the form of Aspergillomarasmin B) in high quantity from axenic cultures of *C. gloeosporioides* on both PDA and OA but it was never found, even in trace amounts, in the olives infected by *Colletotrichum* species irrespective of the cultivar and the *Colletotrichum* species tested, suggesting this toxin is not a key factor in the pathogenesis of OA. Overall, results of this study highlight the diversity and complexity of biochemical interactions between *Colletotrichum* species and olive and stress the importance of *in planta* studies to get a better insight into the pathogenesis mechanisms of OA.

